# Production of a tributyltin-binding protein 2 knockout mutant strain of Japanese medaka, *Oryzias latipes*

**DOI:** 10.1101/2020.04.19.048728

**Authors:** Yoko Kato-Unoki, Yuki Takai, Yosuke Nagano, Satoshi Matsunaga, Shintaro Enoki, Takumi Takamura, Sangwan Kim, Masato Kinoshita, Takeshi Kitano, Yohei Shimasaki, Yuji Oshima

## Abstract

Tributyltin-binding proteins (TBT-bps), members of the lipocalin family, bind TBT in fish blood and are presumed to contribute to detoxification of TBT. Recent studies have shown that many fish species have TBT-bp genes, and that these genes are induced by stresses such as exposure to chemicals or fish pathogenic bacteria. However, the function of TBT-bps, and the mechanisms of their induction and detoxification activity are still unclear. Here, towards elucidating the functions of TBT-bp2, we produced a TBT-bp2 knockout (TBT-bp2^-/-^) strain of Japanese medaka, *Oryzias latipes*, by using the CRISPR/Cas9 system. Gene expression of the mutated TBT-bp2 was reduced, and the cDNA sequencing and predicted protein structure suggested possible loss of function. However, the fish could be grown under normal conditions. Exposure of the TBT-bp2^-/-^ strain of medaka to various stresses in future experiments is expected to contribute to our understanding of this novel detoxification system in aquatic organisms.

## 1. Introduction

Tributyltin-binding proteins (TBT-bps) bind TBT, a typical endocrine disruptor, and are presumed to be involved in detoxification of accumulated TBT (Satone et al., 2008; Shimasaki et al., 2002). As such, they may bind xenobiotics in a novel detoxification pathway. TBT-bps were first discovered in Japanese flounder, *Paralichthys olivaceus* (Shimasaki et al., 2002), but later studies have shown that these proteins and/or genes, and their homologs are widely expressed in many fish species: e.g., tiger puffer, *Takifugu rubripes* (Oba et al., 2011); European eel, *Anguilla anguilla* L. (Nogueira et al., 2009), and Japanese medaka, *Oryzias latipes* (Hashiguchi et al., 2015; Nassef et al., 2011a).

There are two types of TBT-bp, TBT-bp1 (Satone et al., 2008; Shimasaki et al., 2002) and TBT-bp2 (Oba et al., 2007) which are components of pufferfish saxitoxin- and tetrodotoxin-binding proteins (PSTBPs) (Hashiguchi et al., 2015; Yotsu-Yamashita et al., 2018). TBT-bps are presumed to be related to alpha 1-acid glycoprotein (AGP)-like lipocalin protein, which is a major mammalian acute-phase protein that binds to low-molecular-weight basic lipophilic drugs and is involved in the inflammatory response (Fournier et al., 2000; Gutiérrez et al., 2000). Previously, we demonstrated that recombinant TBT-bp1 of Japanese flounder binds to and decreases the cytotoxicity of TBT *in vitro* (Satone et al., 2011). We also reported that TBT-bp2 is highly up-regulated by exposure to TBT in *P. olivaceus* (Nassef et al., 2011b). Response of TBT-bps to chemicals and bacteria has been reported in many fish species: e.g., 5-pentachlorobiphenyl (PCB126) and a PCB mixture (Kanechlor-400, KC-400) in Japanese medaka (Nakayama et al., 2008); 7,12-dimethylbenz [a] anthracene (DMBA) in European eel (Nogueira et al., 2009); and pathogenic bacteria *Edwardsiella tarda* and *Aeromonas hydrophila* in marine medaka (*O. melastigma*) (Dong et al., 2017) and carp (*Labeo rohita*) (Robinson et al., 2014), respectively. Thus, TBT-bps might be stress-responsive proteins that act as acute-phase proteins, the same as mammalian AGP. However, the mechanisms of the induction response and detoxification of TBT by TBT-bps, and its original role (i.e., before man-made chemicals and pathogenic bacteria) remain to be elucidated.

The function of TBT-bps could be clarified by knockout of TBT-bp genes *in vivo*. In recent years, Clustered Regularly Interspaced Short Palindromic Repeats (CRISPR)/CRISPR-associated protein 9 (Cas9) system–based RNA-guided endonucleases have been rapidly developed as a simple and efficient tool for targeted genome editing in a wide range of taxa, including fish (e.g., Ansai and Kinoshita, 2014; Edvardsen et al., 2014; Wiedenheft et al., 2012). Like zinc-finger nucleases and transcription activator-like effector nucleases, this system can efficiently induce sitespecific DNA double-stranded breaks, resulting in targeted gene disruption through indels (insertions or deletions), or targeted gene integration by homologous recombination. Frameshift mutations, which affect the reading frame during translation, are particularly useful for knocking out gene expression or causing gene dysfunction. These genomeediting methods are also used for aquatic toxicology in, for example targeted mutagenesis of aryl hydrocarbon receptor 2a and 2b genes in Atlantic killifish (*Fundulus heteroclitus*) (Aluru et al., 2015) and knockout of multi-resistance associated protein 1 gene in zebrafish (*Danio rerio*) (Tian et al., 2017). Previously, we succeeded in knocking out *T. rubripes* PSTBP2, which has a tandem repeat of domains homologous to TBT-bps (Kato-Unoki et al., 2018). However, no knockout fish strain for TBT-bps has been developed. TBT-bp knockout medaka might will be the best model fish for biology and aquatic toxicology.

To elucidate the role of TBT-bp2 *in vivo*, we knocked out the TBT-bp2 in Japanese medaka, *O. latipes*, by using the CRISPR/Cas9 system, and produced a TBT-bp2^-/-^ (TBT-bp2 KO) strain. We also confirmed the basic function, survival rate, and growth in this strain under normal maintenance conditions.

## 2. Materials and methods

### 2.1 Animals

We used a Japanese medaka strain bred in our laboratory. Medaka adults, embryos, and larvae were maintained under a 14-h/10-h light/dark cycle at 27 ± 1 °C. The fish were fed with *Artemia nauplii* juveniles (24 h incubation) two times per day. Embryos were maintained in Embryo culture medium (ECM: 0.1% NaCl, 0.003% KCl, 0.004% CaCl_2_-H_2_O, 0.008% MgSO_4_, 0.0002% Methylene Blue, pH 7.0) at the above illumination cycle and temperature. The use of animals in this study, and our genome-editing procedures, were in accordance with Kyushu University’s guidelines for genetic recombination and animal experimentation. The TBT-bp2 KO strain produced in this study was deposited in the Material Management Center (https://mmc-u.jp/; Material number QM2017-0041) of Kyushu University, Japan.

### 2.2 Selection of the target site

TBT-bp2–homologous genes were found in the medaka database (NIG/UT MEDAKA1/oryLat2 (Oct. 2005)) in the Genome Browser Gateway (http://genome-asia.ucsc.edu/cgi-bin/hgGateway?redirect=manual&source=genome.ucsc.edu) by BLAT Search Genome with TBT-bp2 cDNA sequences (GenBank accession nos. LC229802–LC229804). Then multiple sequence alignments were conducted using the program MAFFT (https://mafft.cbrc.jp/alignment/server/). The target exon and TBT-bp2–specific target site for CRISPR/Cas9 were selected by using the target design web site, CRISPRdirect (https://crispr.dbcls.jp) (using the same database in the Genome Browser Gateway).

### 2.3 Preparation of single-guide RNA and Cas9 mRNA

Single-guide RNA (sgRNA) for the target sequence and Cas9 mRNA were prepared by using the pDR274 vector (plasmid 42250, Addgene, Watertown, MA, USA) and pCS2+hSpCas9 vector (plasmid 51815, Addgene), in accordance with published methods (Ansai and Kinoshita, 2014; Kato-Unoki et al., 2018). The annealed oligonucleotide sequences for the sgRNA were as follows (underlined letters indicate a cohesive sequence with the vector): sgRNA sense segment, 5’-<u>TAGG</u>ACAACGAGAGGAACGTGTAC; sgRNA anti-sense segment, 5’-<u>AAAC</u>GTACACGTTCCTCTCGTTGT.

### 2.4 Microinjection

Yamamoto’s Ringer’s solution (0.75% NaCl, 0.02% KCl, 0.02% CaCl_2_, 0.002% NaHCO_3_, pH 7.3) containing 100 ng/μL of Cas9 mRNA and 25 ng/μL of sgRNA was injected into Japanese medaka 39 embryos at the one-cell stage using Femto Jet (Eppendorf, Hamburg, Germany) and GD-1 needles (Narishige, Tokyo, Japan) in accordance with published methods (Ansai and Kinoshita, 2014). Control embryos, 30 individuals, were only injected with Yamamoto’s Ringer’s solution.

### 2.5 Genomic DNA extraction

At the first day post fertilization (dpf), the egg envelope was broken with forceps in accordance with published methods (Ansai and Kinoshita, 2014), and embryos were lysed individually in 25 μL of alkaline lysis solution containing 25 mM NaOH and 0.2 mM EDTA and incubated at 95 °C for 15 min. Each sample was neutralized with 25 μL of 40 mM Tris-HCl (pH 8.0) and used as a genomic DNA sample. To determine the mutation pattern in adult fish, genomic DNA was extracted from fin and/or egg as above.

### 2.6 Heteroduplex mobility assay and genotyping

To identify mutations induced by sgRNA in the genomic target site, we performed the heteroduplex mobility assay (HMA) as described previously (Ansai and Kinoshita, 2014). Briefly, the genomic region containing the target sequence of the sgRNA was amplified by polymerase chain reaction (PCR) using KOD-FX DNA polymerase (Toyobo, Osaka, Japan) with primers HMA-F and HMA-R (Table 1 and Fig. 1). The PCR mixture contained 1 μL of genomic DNA, 1× PCR buffer for KOD FX, 0.4 mM of each dNTP, 0.2 μM of each primer, and 0.05 units of KOD FX DNA polymerase in a total volume of 10 μL. The PCR conditions were one cycle at 94 °C for 2 min followed by 35 cycles of 98 °C for 10 s, 60 °C for 20 s, and 68 °C for 20 s. The resulting amplicons were analyzed by 10%T, 5%C polyacrylamide gel electrophoresis in Tris/borate/EDTA buffer.

**Table 1.**
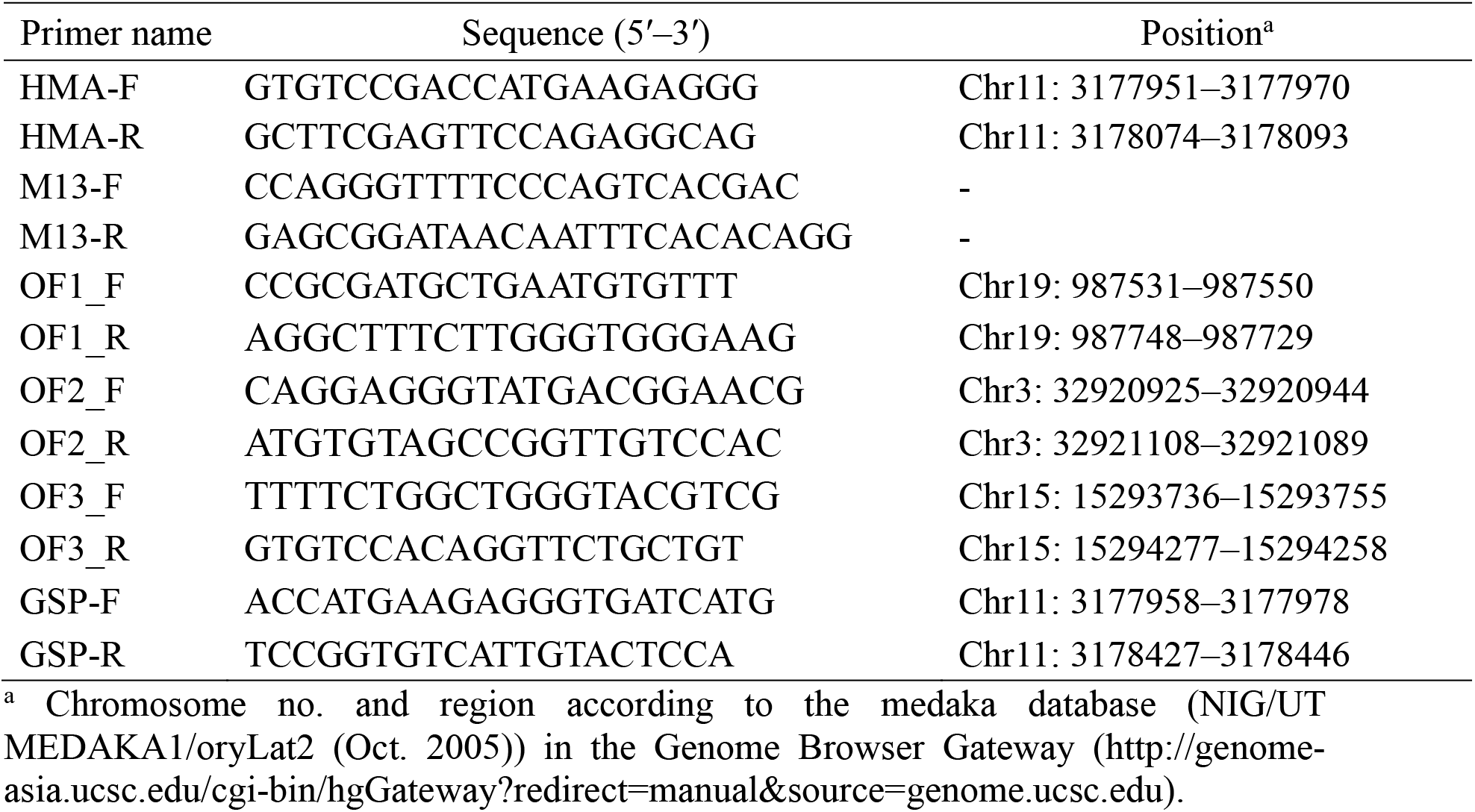
Primers used in this study. Primers were designed by using Primer-BLAST (https://www.ncbi.nlm.nih.gov/tools/primer-blast/)

**Fig. 1.**
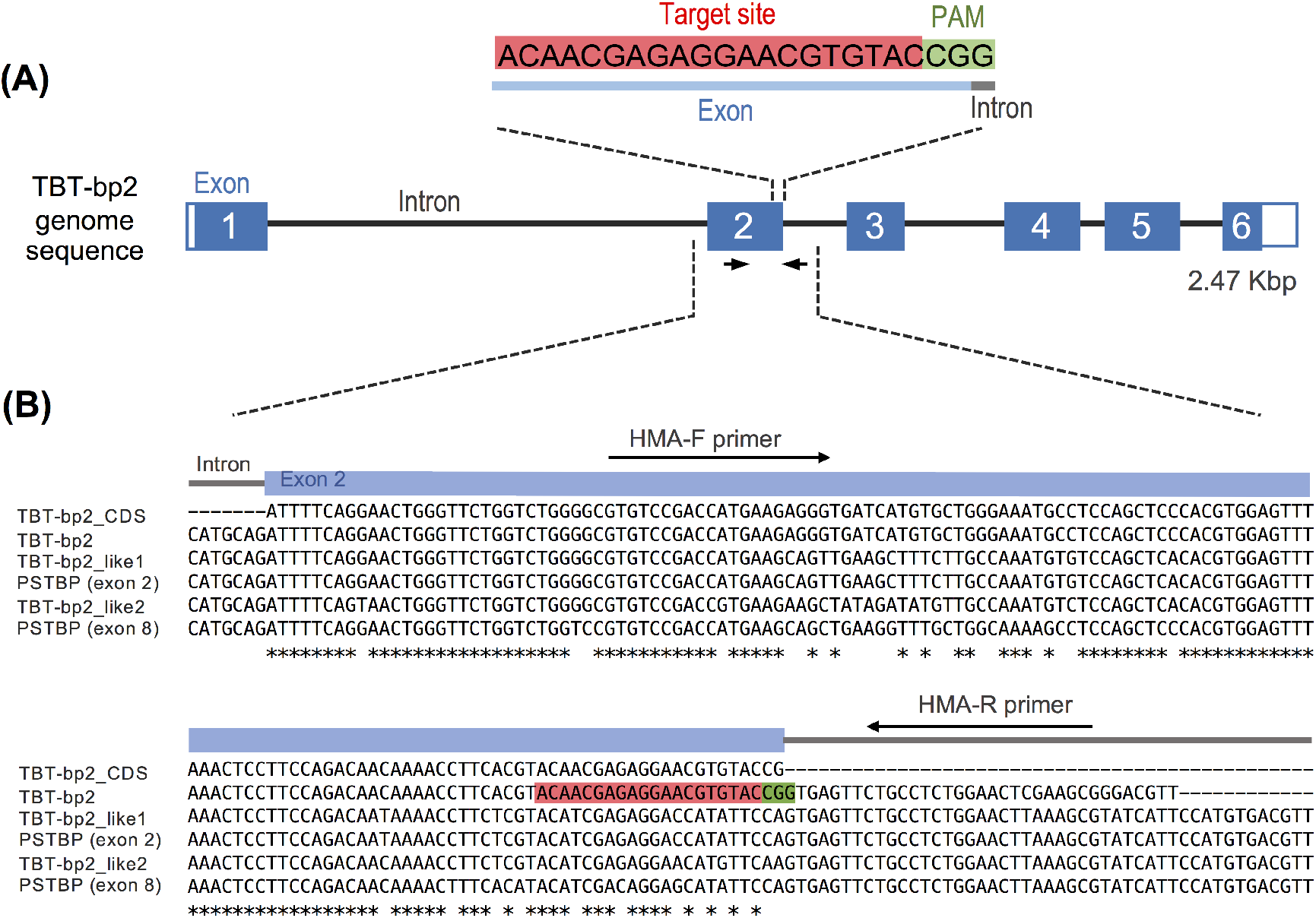
Schematic of the TBT-bp2 target site. A) Genomic sequence of *Oryzias latipes* TBT-bp2 and the target site. Blue boxes and black bars indicate exons and introns, respectively. Red and green boxes are target and PAM (protospacer adjacent motif) sequences, respectively, for CRISPR/Cas9. Black arrows show the positions of HMA primers (see Table 1). B) Multi-alignment of the cDNA and genomic sequences of TBT-bp2 (TBT-bp2_CDS and TBT-bp2, respectively), and the sequences of homologous genes (TBT-bp2_like1, TBT-bp2_like2, PSTBP (exon 2), and PSTBP (exon 8); the sequences were obtained from the medaka database (NIG/UT MEDAKA1/oryLat2 (Oct. 2005)) in the Genome Browser Gateway (http://genome-asia.ucsc.edu/cgi-bin/hgGateway?redirect=manual&source=genome.ucsc.edu) as follows [gene name, chromosome number: alignment region]: TBT-bp2, chr11: 3177914–3178101; TBT-bp2_like1, chr11: 3255415–3255614; TBT-bp2_like2, chr11: 3236109–3236308 (complement); PSTBP (exon 2), chr11: 3222335–3222534 (complement); PSTBP (exon 8), chr11: 3211331–3211530 (complement); TBT-bp2_CDS, GenBank accession no. LC229802. Designations are as in panel A. Dashes and asterisks indicate gaps and conserved nucleotide in all six sequences, respectively.

Genotyping for TBT-bp2 mutations was performed by using the same primers and PCR conditions as described for the HMA.

### 2.7 Sequence analysis

To confirm the mutations, PCR products in the HMA were treated with 10×A-attachment mix (Toyobo), and then subcloned into a pTac-1 vector (BioDynamics Laboratory Inc., Tokyo, Japan). The fragments containing the cloned genomic sequence were amplified from each colony by using the primers M13-F and M13-R (Table 1), and then sequenced with the M13-F primer by Eurofins Genomics Ltd. (Tokyo, Japan). Direct sequencing of a PCR fragment was performed by using one side primer of that PCR primer.

### 2.8 Production of the TBT-bp2 KO strain

Injected F0 embryos (27 individuals) remaining after HMA were reared to produce the TBT-bp2 KO strain. F0 mutants harboring a 7-nucleotide deletion were selected and naturally mated with wild-type (WT) medaka in a tank. The genotypes of F1 individuals were determined from fin samples by using HMA primers after the fish reached the adult stage, and male and female F1 heterozygous (-/+) mutants for the 7-nucleotide deletion were naturally mated with each other. The resultant homozygous F2 mutants (-/-), TBT-bp2 KO, were reared for further experiments.

### 2.9 cDNA sequencing, gene expression and protein structure prediction

To determine the TBT-bp2 KO cDNA sequence, total RNA of the TBT-bp2 KO strain was extracted by using ISOGEN II (Nippon Gene, Tokyo, Japan), and mRNA sequencing (mRNA-Seq) by next-generation sequencing was performed by Chemical Dojin Co. (Kumamoto, Japan). The quality of reads in the FASTQ file was checked by using FastQC (ver.0.11.9, https://www.bioinformatics.babraham.ac.uk/projects/fastqc/). Short fragments and low-quality sections of mRNA reads were removed by using Trimmomatic (ver.0.39, http://www.usadellab.org/cms/index.php?page=trimmomatic). Non-chromosomal mRNA reads (ribosomal and mitochondrial mRNA reads) were removed by using SortMeRNA (ver.2.1, https://bioinfo.lifl.fr/RNA/sortmerna/). The cleaned mRNA reads were analyzed with STAR (ver.2.7.3a, https://github.com/alexdobin/STAR), SAMtools (ver.1.10, http://www.htslib.org), featureCounts (ver.2.0.0, http://subread.sourceforge.net/), and edgeR (ver.3.26.8, https://bioconductor.org/packages/release/bioc/html/edgeR.html). The genome sequence and annotation information of Japanese medaka were downloaded from Ensembl (ver.ASM223467v1.99, http://asia.ensembl.org/index.html). Gene expression was quantified as transcripts per kilobase million (TPM) (Wagner et al., 2012). Significant difference between WT and KO were detected by Fisher’s exact test. P-values less than 0.01 were classed as statistically significant.

The protein structure of TBT-bp2 in the KO strain was predicted from the above cDNA sequence with the homology modeling computer program, Protein Homology/analogY Recognition Engine V 2.0 (Phyre2) (http://www.sbg.bio.ic.ac.uk/phyre2/html/page.cgi?id=index).

### 2.10 Off-target analysis

The protospacer adjacent motif (PAM) (Streptococcus pyogenes based: 5’-NGG) is the recognition sequence of Cas9 (Mojica et al., 2009), and a 10–12 base sequence in the PAM-proximal region of the target sequence is reported to be important for mediating Cas9 binding (Cho et al., 2014; Kuscu et al., 2014). Hence, to search for off-target candidates by using the above target design web site, up to 2 mismatches were allowed in the target sequence plus the PAM (20 mer+PAM) and complete matches were required in the 12-mer targeting sequence followed by a PAM (12 mer+PAM). Each off-target region was PCR amplified from fin genomic DNA of the KO strain with each primer set (Table 1). The PCR mixture and conditions were as follows: 0.75 μL genomic DNA, 1× Phusion HF buffer, 0.2 mM of each dNTP, 0.2 μM of each primer, and 0.1 μL Phusion DNA polymerase in a total volume of 10 μL, with one cycle at 98 °C for 2 min, followed by 40 cycles of 98 °C for 10 s, 65 °C (primer OF3_F and OF3_R: 63 °C) for 20 s, and 72 °C for 30 s. The PCR products were directly sequenced.

### 2.11 Analysis of growth

From the hatch, three genotypes, WT, heterozygotes (-/+) and homozygotes (-/-; KO), were reared in three separate tanks with 20 individuals per tank. All tanks were maintained under the same conditions throughout the 3 months of experiments. Five fish were selected randomly per tank every month, anesthetized with FA100 (Tanabe Pharmaceuticals, Osaka, Japan), wiped to remove water, and individual body weight and length were measured. Significant difference between control and each genotype were detected by Welch’s *t*-test, respectively. *P*-values less than 0.05 were classed as statistically significant.

## 3. Results

### 3.1 Selection of the target site

Partial sequence alignments of the four TBT-bp2–homologous genes detected in the medaka database are shown in Fig. 1: [gene name in this study, chromosome number: alignment region] TBT-bp2_like1, chr11: 3255098–3255614; TBT-bp2_like2, chr11: 3236109–3236634 (complement); PSTBP (exon 2), chr11: 3222335–3222859 (complement); and PSTBP (exon 8), chr11: 3211331–3211855 (complement). These genes correspond to those in Hashiguchi et al. (2015) as follows: [gene name in this study / gene name in Hashiguchi et al. (2015)] TBT-bp2_like1 / Olat_TBT_bp2_like1; TBT-bp2_like2 / Olat_TBT_bp2_like2; PSTBP (exon 2) and PSTBP (exon 8) / Olat_PSTBP. From this alignment, we identified a 20-nucleotide region with sufficiently low similarity to be a specific target site, 5’-ACAACGAGAGGAACGTGTAC (red box in Fig. 1) at the 3’ end of exon 2. The PAM sequence was only present in the TBT-bp2 gene among the aligned sequences.

### 3.2 Confirmation of the mutation induced by TBT-bp2 sgRNA

One day after the injection of TBT-bp2 sgRNA and Cas9 mRNA, hetero-duplexed bands, which indicate the occurrence of mutations in an HMA analysis, were observed in 7 embryos (embryo nos. 3, 5, 8–12) out of 12 randomly selected embryos (mutation rate, 58.3%) (Fig. 2). The survival rate of TBT-bp2 sgRNA–treated embryos at 1 day after injection was 30/39 (76.9%), whereas that of the control was 27/30 (90.0%). Although the survival rate of the TBT-bp2 sgRNA–treated embryos was lower than that of the control, there was no significant difference.

**Fig. 2.**
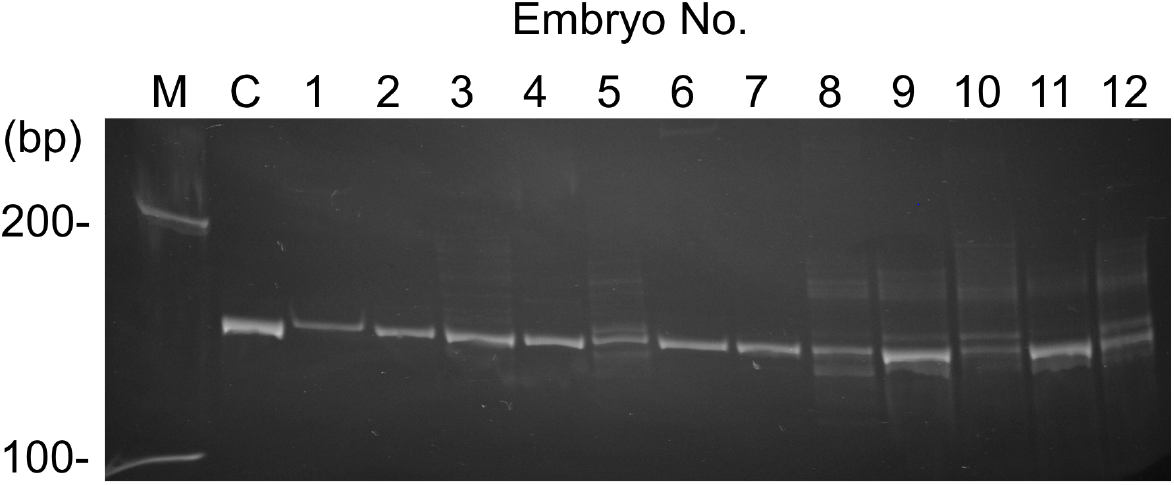
Heteroduplex mobility assay (HMA) of embryos injected with TBT-bp2 sgRNA. Lane M is DNA marker and Lane C is a control sample injected only with Yamamoto’s Ringer’s solution. Lanes 1 through 12 correspond to embryo numbers.

The mutations observed in the target sequences were confirmed by sequencing. PCR products taken from the heteroduplex bands for embryos no. 8 and 10 were subcloned and the mutated sequences were determined (Fig. 3). The rates of mutated sequences induced in individual embryos were low (embryo no. 8, 4/29 sequences (23.8%); embryo no. 10, 5/20 sequences (25.0%)); however, the frameshift mutations that knocked out TBT-bp2 (mutation types, −2, −7, −8 in Fig. 3) occurred at high rates in the mutated sequences: [embryo no., frameshift mutation sequences/mutation sequences (%)] embryo no. 8, 3/4 (75%); embryo no. 10, 4/5 (80%). In addition, most of the mutations were considered to be induced via microhomology-mediated end joining (MMEJ; underlined letters in Fig. 3; McVey and Lee, 2008).

**Fig. 3.**
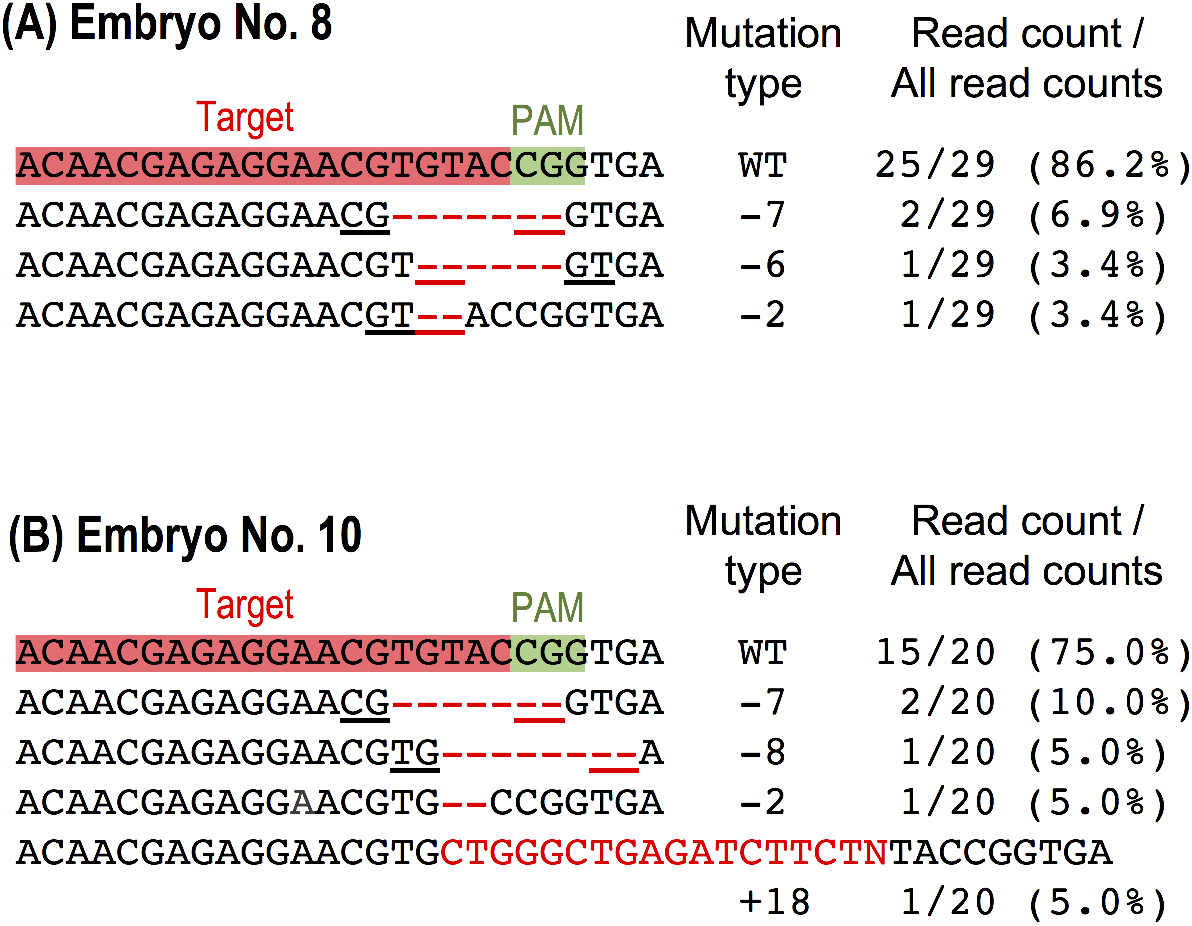
Mutations induced by TBT-bp2 sgRNA. Subcloned sequences of the target locus observed in embryos no. 8 (A) and no. 10 (B) injected with TBT-bp2 sgRNA. Red and green boxes are the sgRNA target and PAM sequences, respectively. Red dashes and letters indicate the identified mutations (deletion and insertion and/or nucleotide substitution). Underlining indicates microhomologies. The number to the right of each sequence indicates the number of mutated nucleotides (-: deletion; +: insertion). The rates show the frequency of detection of the mutated sequence among total sequences.

### 3.3 Production of the TBT-bp2 KO strain

Homozygotes (-/-; KO) were produced in the F2 generation. As shown in Supplementary Table S1, the mutant rates in the F1 generation were 37.5% for F1-1 and 27.3% for F1-2 (F1-1 and F1-2 were individuals from one fertilization). Several individuals of both sexes that were heterozygous for a 7-nucleotide deletion were obtained, enabling the production of homozygotes by mating. The proportions of the three genotypes in randomly selected 48 F2 embryos were as follows (Supplementary Table S1): WT, 18.8%; heterozygotes (-/+), 64.6%; and homozygotes (-/-; KO), 16.7%.

### 3.4 Off-target analysis

We found three possible off-target sites (named OF1, 2, 3) for the TBT-bp2 sg RNA. One off-target site (OF1; two mismatches) was found in a 20 mer +PAM, and two off-target sites (OF2 and OF3) were found in a 12 mer + PAM (Table 2 and Supplementary Fig. S1). OF1 and OF2 were in the non-coding region, whereas OF3 might be a coding site since it is found in EST (expressed sequence tag) sequences of that region in the database. No mutation was detected in these sites in the KO strain by PCR amplification of sequences of 3 individuals (Supplementary Fig. S1).

**Table 2.**
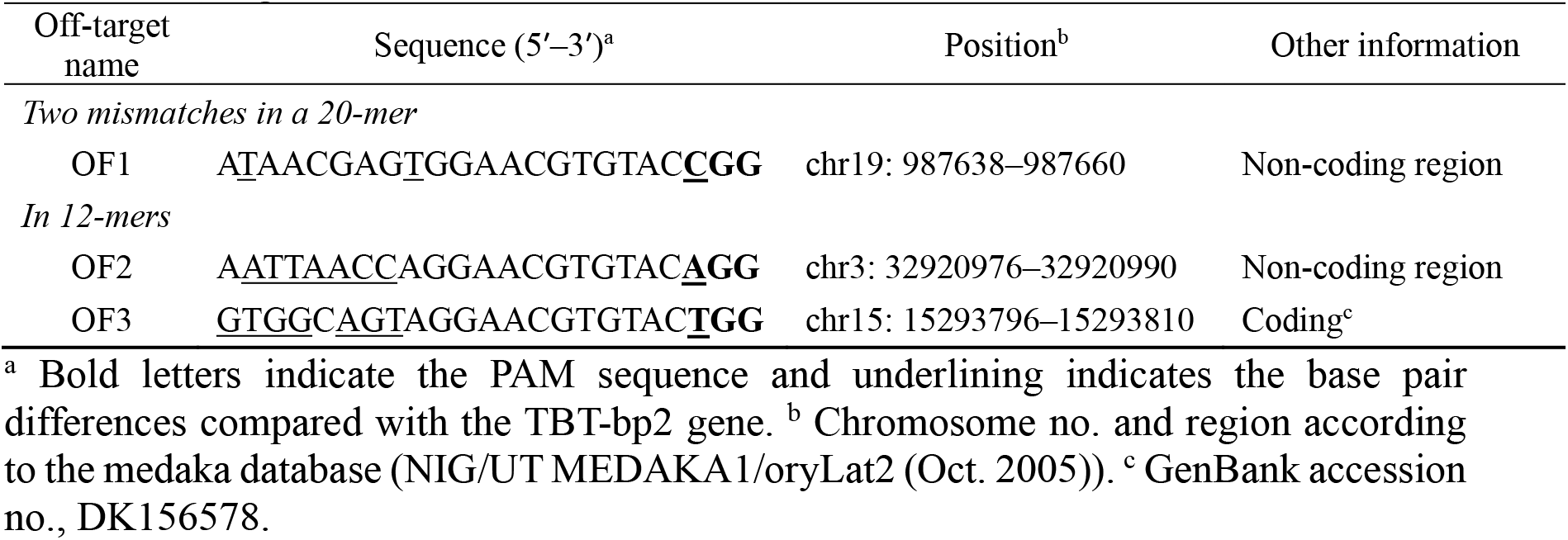
Off-target list.

### 3.5 cDNA sequencing, gene expression and protein structure prediction

The RIN (RNA integrity number) of the extracted total RNA of each of the 4 WT and KO individuals was higher than 8.0. From the TBT-bp2 mRNA-Seq read sequences, we confirmed that the TBT-bp2 cDNA sequence in the KO strain contained the 7-nucleotide deletion (Fig. 4A). The deduced amino acid sequence showed a frame shift starting in exon 2 and a premature stop codon at position 99 from the N-terminus. From the result of the Phyre2 analysis, the predicted KO protein structure was quite different from that of WT, and the truncated protein may not form the barrel-like structure that seems to be functionally important for ligand binding in lipocalin protein family (Fig. 4B; Flower, 1996; Satone et al., 2008).

**Fig. 4.**
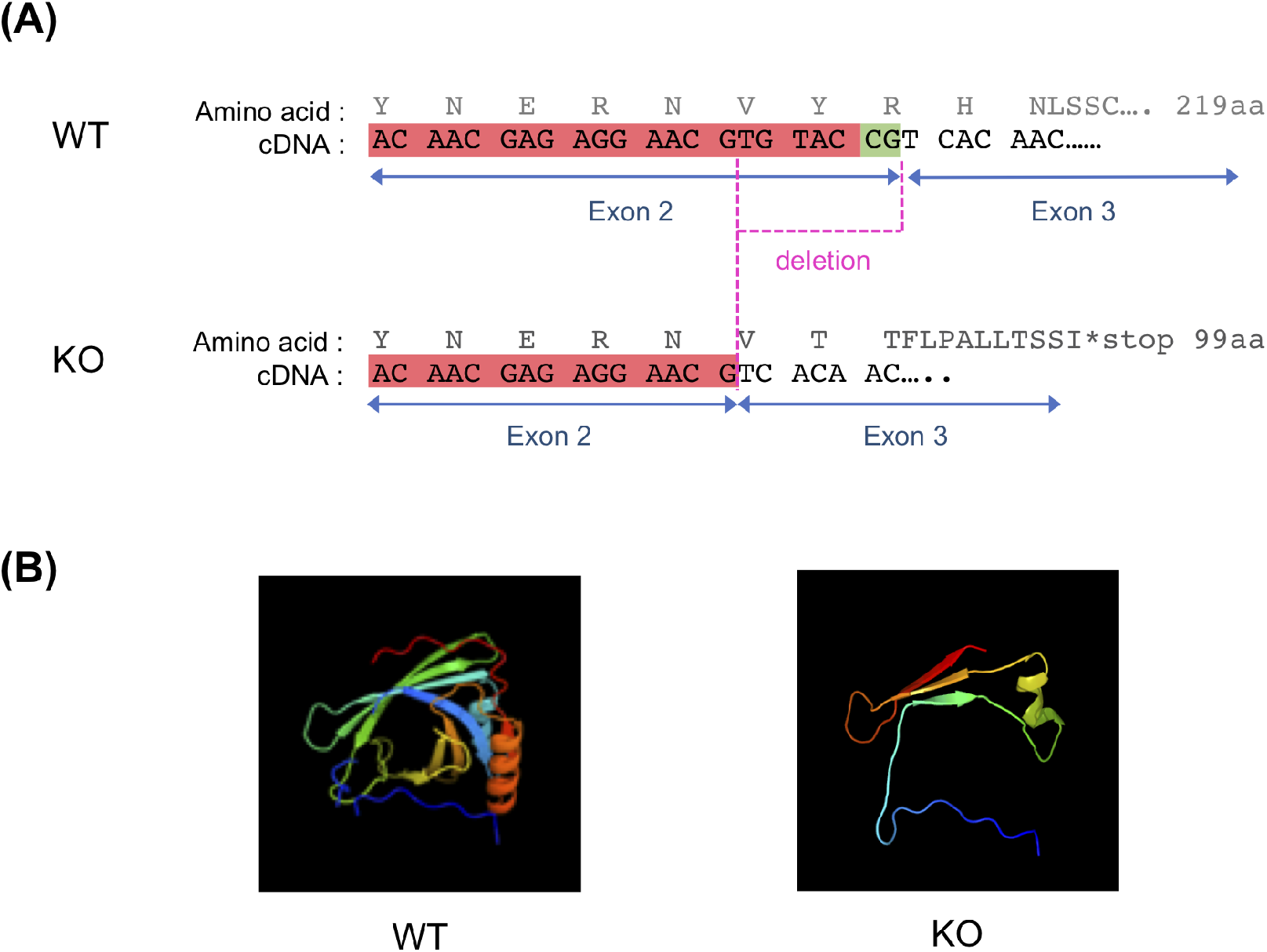
TBT-bp2 cDNA sequence and predicted protein structure of TBT-bp2 in the WT and KO strains. A) The TBT-bp2 cDNA and deduced amino acid sequences determined by mRNA-Seq. Red and green bars are target and PAM sequences for CRISPR/Cas9, respectively. B) Predicted TBT-bp2 protein structures for the WT and KO strains. The structures were predicted by Protein Homology/analogY Recognition Engine V 2.0 (Phyre2) using the cDNA sequence determined by mRNA-Seq. Each structure is the top model based on the template, c2wewA (http://www.sbg.bio.ic.ac.uk/phyre2/html/flibview.cgi?pdb=c2wewA). Individual β-strands and α-helices are indicated by arrows and springs, respectively.

The expression of TBT-bp2 was significantly reduced in the KO strain (0.8 TPM) compared with WT (42.5 TPM) (logFC −5.8, *P* < 0.01 Fisher’s exact test), whereas there was no significant difference in β-actin between KO and WT (logFC −0.34, *P* = 0.32) (Fig. 5A). Thus, the results of cDNA sequencing and prediction of the protein structure suggest loss of function of TBT-bp2 in the KO strain.

**Fig. 5.**
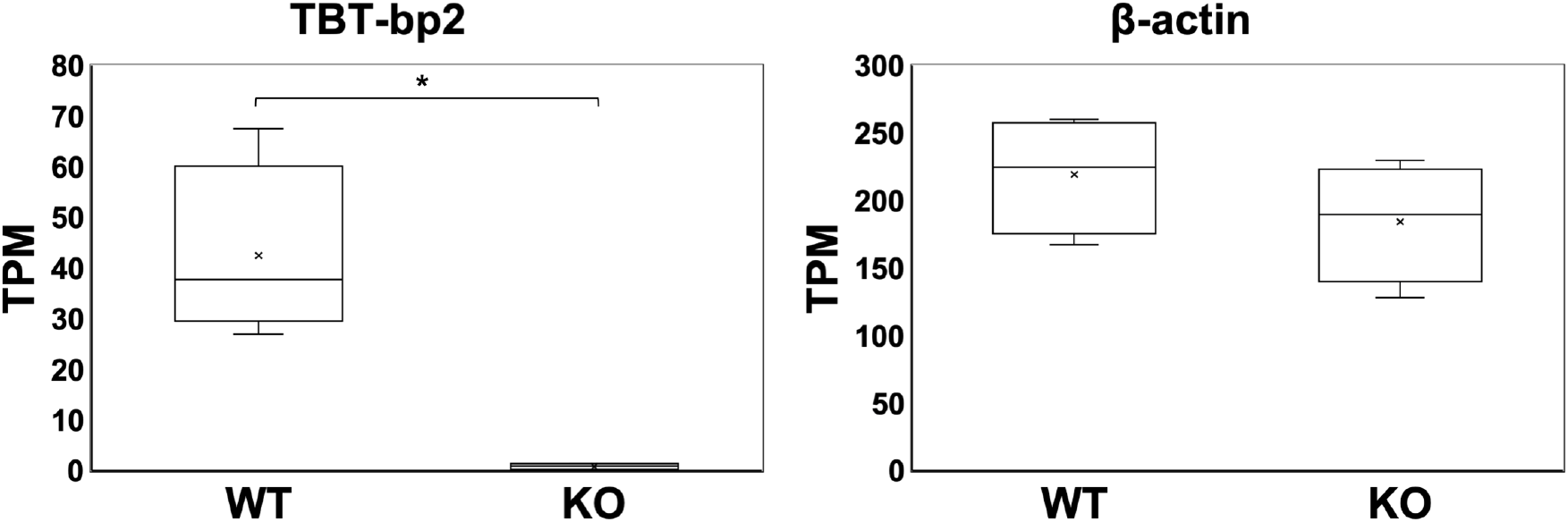
Expression of the TBT-bp2 gene in WT and KO strains. Boxplots of the expression levels of TBT-bp2 and β-actin genes in 4 WT and 4 KO individuals. The expression levels of each gene were quantified as transcripts per kilobase million (TPM) from the mRNA-Seq data. The internal line shows the median. * *P* < 0.01 (significant difference, Fisher’s exact test).

### 3.6 Basic function of TBT-bp2 KO strain in normal conditions

Measurements of growth (body weight and length) among the genotypes (WT, -/+, and -/-) were not significantly different from each other at 3 months after hatching (Fig. 6). No significant differences in body weight between genotypes were observed at 1 month after hatching (Fig. 6A). Body weight at 2 months after hatching was significantly higher in the heterozygotes (-/+) than in WT (Fig. 6C), but this difference disappeared by 3 months after hatching (Fig. 6E). Body length was significantly lower in homozygotes (-/-) than in WT at 1 month after hatching (Fig. 6B); however, at 2 months after hatching, body length was significantly higher in heterozygotes (-/+) than in WT and there was no longer a significant difference between homozygotes and WT (Fig. 6D). No significant differences in body length were detected at 3 months (Fig. 6F). We conclude that there was no obvious effect of the KO (-/-) on growth.

**Fig. 6.**
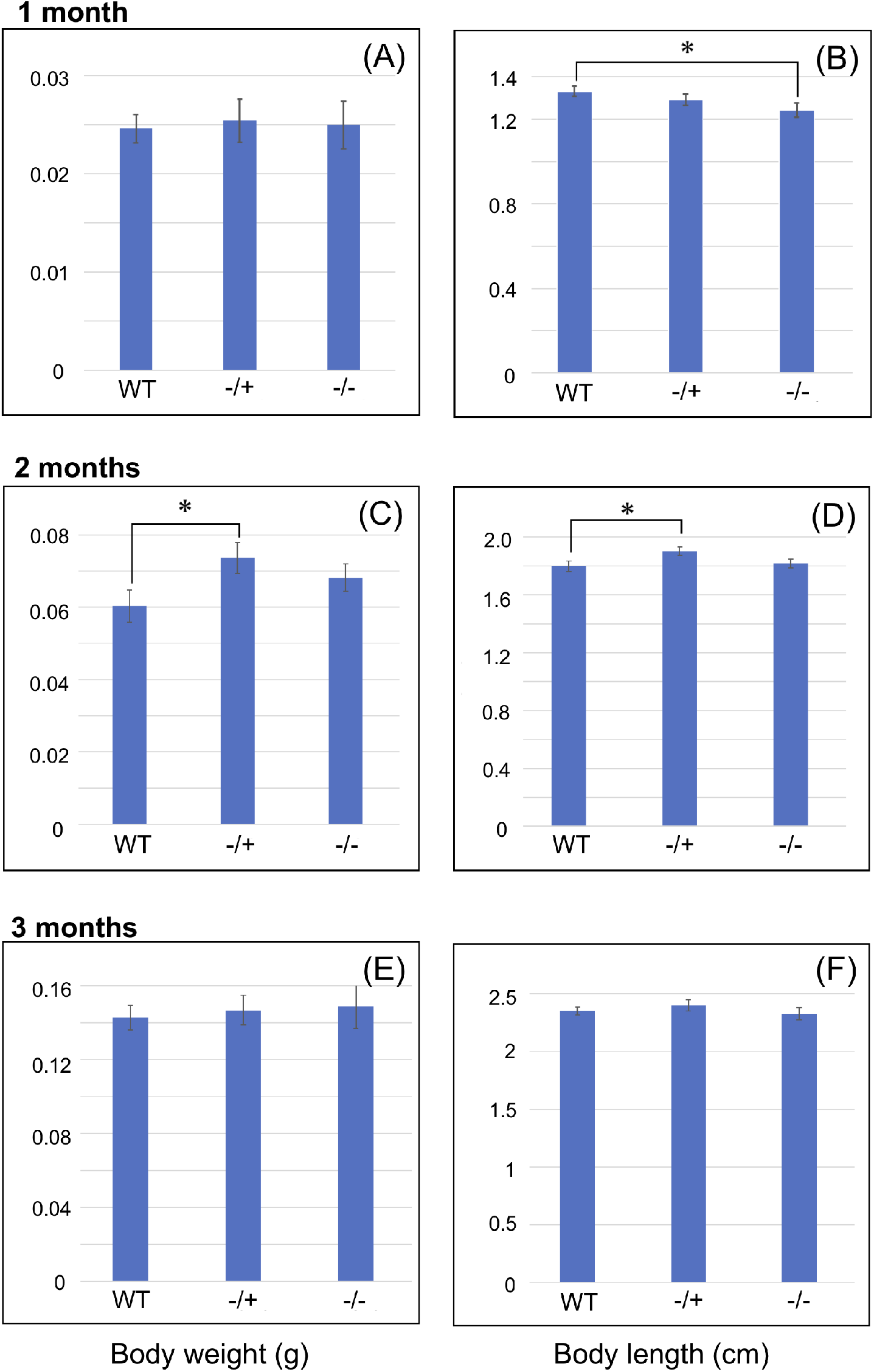
Comparison of growth among genotypes. Measures of growth at 1 month (A, B), 2 months (C, D), and 3 months (E, F) after hatching are displayed. A), C), E), Body weight; B), D), F), body length. WT, wild-type; -/+, heterozygote; -/-, homozygote (i.e., KO strain). Values are means ± SD (n = 15 from 3 tanks, 5 fishes per tank). * *P* < 0.05 (significant difference, Welch’s *t*-test).

## 4. Discussion

Here we report the development of a strain of Japanese medaka with the TBT-bp2 gene specifically knocked out by the CRISPR/Cas9 system. To avoid off-target mutations in the TBT-bp1 gene and homologous genes (Hashiguchi et al., 2015), we used the medaka database and multiple sequence alignment to select a TBT-bp2 gene target region with low similarity to the other genes. In addition, the CRISPR/Cas9 system requires a PAM sequence (Mojica et al., 2009). Using the 20-nucleotide TBT-bp-specific gene target sequence and the PAM sequence, we designed the TBT-bp2–specific sgRNA and successfully knocked out the TBT-bp2 gene (Fig. 1). No mutation was found in the 3 possible off-target sites (OF1–OF3) in the KO strain (Supplementary Fig. S1).

This study demonstrates that producing a KO strain is possible even with a low mutation rate. As mentioned in section 3.2, the individual mutation rates of the two F0 embryos examined were low (23.8% and 25.0%; Fig. 3). Of these, the rates of the 7-nucleotide deletion were only 6.9% and 10%, respectively. Nevertheless, we obtained both male and female F1 heterozygotes for the 7-nucleotide deletion from the F0 founder (Supplementary Table S1). In a previous report, the variation and frequencies of mutation types produced by the CRISPR/Cas9 system in red sea bream (*Pagrus major*) differed among tissues (Kishimoto et al., 2018). Our result for F1 mutants supports this previous report (F1-1 and F1-2 in Supplementary Table S1).

The TBT-bp2 KO medaka strain produced here might be a useful model fish for ecotoxicological studies—not only for TBT-bp2 function analysis but also for understanding the novel detoxification pathway. The expression of TBT-bp2 was significantly reduced in the KO strain compared with WT (Fig. 5), and from the predicted protein structure, the TBT-bp2 KO medaka probably lost its ability to bind TBT (Fig. 4B), whereas no consistent differences were detected in survival rate or growth of TBT-bp2 KO medaka compared with WT (Section 3.2 and Fig. 6). For functional analyses, e.g., toxicant exposure experiments, it is important that a KO animal has no abnormalities except target function. As mentioned in the Introduction, TBT-bp2 is presumed to bind, accumulate and detoxify low-molecular lipophilic compounds such as TBT (e.g., Nakayama et al., 2008; Satone et al., 2008; Shimasaki et al., 2002). Therefore, we hypothesize that reduced or no accumulation and/or detoxification of TBT occurs in the TBT-bp2 KO medaka. As recombinant PSTBP1 and 2 of *T. rubripes*, which is a duplicate-fused type of TBT-bp2, can bind tetrodotoxin (TTX) and TBT (Satone et al., 2017), this hypothesis on the function of TBT-bp2 could be tested by chemical exposure of the TBT-bp2 KO medaka.

## Acknowledgment

This work was funded by the Japan Society for the Promotion of Science (JSPS) KAKENHI grant number JP18H02280.

---

**Supplementary Table S1.** Rates of genotypes in the F1 and F2 generations

| Genotype | Number<sup>a</sup> (%) |
| --- | --- |
| ***F1-1*<sup>b</sup>** |  |
| WT | 5/8 (62.5) |
| 7-nucleotide deletion (heterozygote: -/+) | 1/8 (12.5) |
| 6-nucleotide deletion (heterozygote: -/+) | 1/8 (12.5) |
| 2-nucleotide insertion and 2 nucleotide substitution (heterozygote: -/+) | 1/8 (12.5) |
| ***F1-2*<sup>b</sup>** |  |
| WT | 8/11 (72.7) |
| 7-nucleotide deletion (heterozygote: -/+) | 2/11 (18.2) |
| 2-nucleotide insertion and 1 nucleotide substitution (heterozygote: -/+) | 1/11 (9.1) |
| ***F2*** |  |
| WT | 9/48 (18.8) |
| 7-nucleotide deletion (heterozygote: -/+) | 31/48 (64.6) |
| 7-nucleotide deletion (homozygote (KO): -/-) | 8/48 (16.7) |

<sup>a</sup> Rates are the number of individuals carrying the genotype/the total number of individuals WT, Wild type. <sup>b</sup> F1-1 and F1-2 are individuals from one fertilization, respectively.

**Supplementary Fig. S1.**
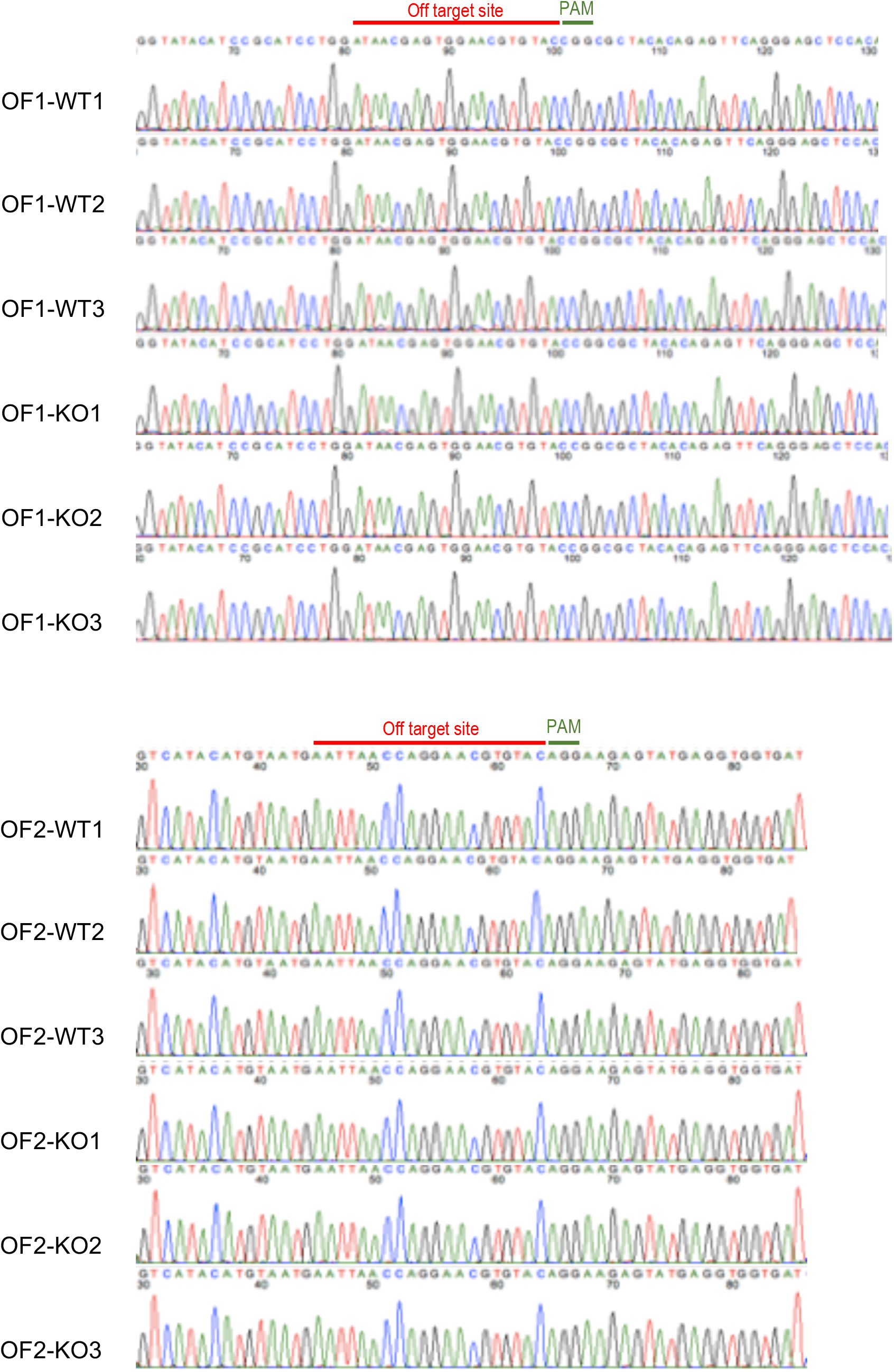
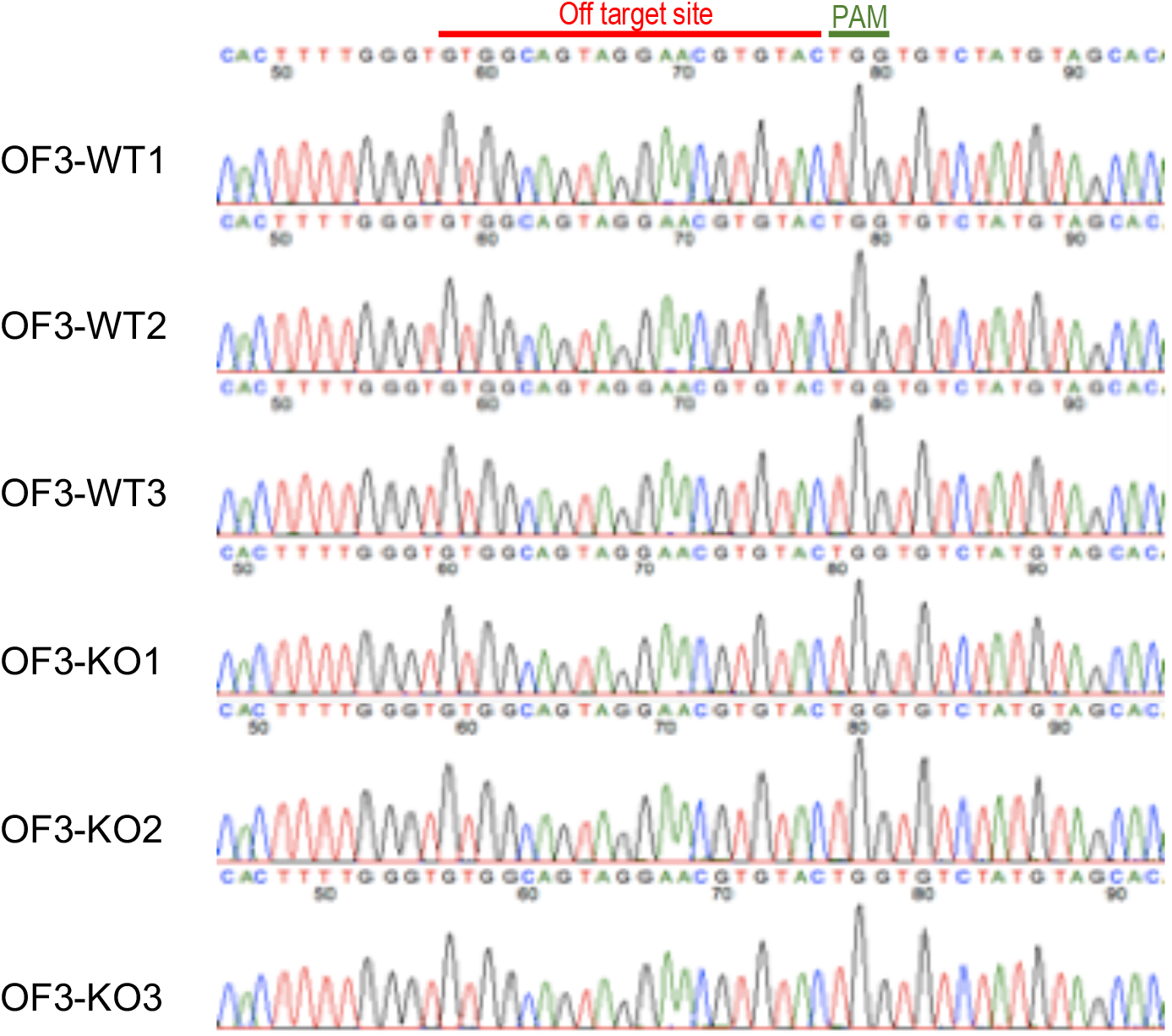
Off-target sequences in WT and KO (n = 3). OF1–OF3 are candidate off-target sequences for the TBT-bp2 sg RNA (see Table 2).

## References

Aluru, N., Karchner, S.I., Franks, D.G., Nacci, D., Champlin, D., Hahn, M.E., 2015. Targeted mutagenesis of aryl hydrocarbon receptor 2a and 2b genes in Atlantic killifish (*Fundulus heteroclitus*). Aquat. Toxicol. 158, 192–201. https://doi.org/10.1016/j.aquatox.2014.11.016

Ansai, S., Kinoshita, M., 2014. Targeted mutagenesis using CRISPR/Cas system in medaka. Biol. Open 3, 362–371. https://doi.org/10.1242/bio.20148177

Cho, S.W., Kim, S., Kim, Y., Kweon, J., Kim, H.S., Bae, S., Kim, J.S., 2014. Analysis of off-target effects of CRISPR/Cas-derived RNA-guided endonucleases and nickases. Genome Res. 24, 132–141. https://doi.org/10.1101/gr.162339.113

Dong, M., Seemann, F., Humble, J.L., Liang, Y., Peterson, D.R., Ye, R., Ren, H., Kim, H.-S., Lee, J.-S., Au, D.W.T., Lam, Y.W., 2017. Modification of the plasma complement protein profile by exogenous estrogens is indicative of a compromised immune competence in marine medaka (*Oryzias melastigma*). Fish Shellfish Immunol. 70, 260–269. https://doi.org/10.1016/j.fsi.2017.09.020

Edvardsen, R.B., Leininger, S., Kleppe, L., Skaftnesmo, K.O., Wargelius, A., 2014. Targeted mutagenesis in Atlantic salmon (*Salmo salar* L.) using the CRISPR/Cas9 system induces complete knockout individuals in the F0 generation. PLoS One 9, e108622. https://doi.org/10.1371/journal.pone.0108622

Flower, D.R., 1996. The lipocalin protein family: Structure and function. Biochem. J. 318, 1–14. https://doi.org/10.1042/bj3180001

Fournier, T., Medjoubi-N, N., Porquet, D., 2000. Alpha-1-acid glycoprotein. Biochim. Biophys. Acta - Protein Struct. Mol. Enzymol. 1482, 157–171. https://doi.org/10.1016/S0167-4838(00)00153-9

Gutiérrez, G., Ganfornina, M.D., Sánchez, D., 2000. Evolution of the lipocalin family as inferred from a protein sequence phylogeny. Biochim. Biophys. Acta - Protein Struct. Mol. Enzymol. 1482, 35–45. https://doi.org/10.1016/S0167-4838(00)00151-5

Hashiguchi, Y., Lee, J.M., Shiraishi, M., Komatsu, S., Miki, S., Shimasaki, Y., Mochioka, N., Kusakabe, T., Oshima, Y., 2015. Characterization and evolutionary analysis of tributyltin-binding protein and pufferfish saxitoxin and tetrodotoxin-binding protein genes in toxic and nontoxic pufferfishes. J. Evol. Biol. 28, 1103–1118. https://doi.org/10.1111/jeb.12634

Kato-Unoki, Y., Takai, Y., Kinoshita, M., Mochizuki, T., Tatsuno, R., Shimasaki, Y., Oshima, Y., 2018. Genome editing of pufferfish saxitoxin-and tetrodotoxin-binding protein type 2 in *Takifugu rubripes*. Toxicon 153, 58–61. https://doi.org/10.1016/j.toxicon.2018.08.001

Kishimoto, K., Washio, Y., Yoshiura, Y., Toyoda, A., Ueno, T., Fukuyama, H., Kato, K., Kinoshita, M., 2018. Production of a breed of red sea bream *Pagrus major* with an increase of skeletal muscle mass and reduced body length by genome editing with CRISPR / Cas9. Aquaculture 495, 415–427. https://doi.org/10.1016/j.aquaculture.2018.05.055

Kuscu, C., Arslan, S., Singh, R., Thorpe, J., Adli, M., 2014. Genome-wide analysis reveals characteristics of off-target sites bound by the Cas9 endonuclease. Nat Biotech 32, 677–683. https://doi.org/10.1038/nbt.2916

McVey, M., Lee, S.E., 2008. MMEJ repair of double-strand breaks (director’s cut): deleted sequences and alternative endings. Trends Genet. 24, 529–538. https://doi.org/10.1016/j.tig.2008.08.007

Mojica, F.J.M., Díez-Villaseñor, C., García-Martínez, J., Almendros, C., 2009. Short motif sequences determine the targets of the prokaryotic CRISPR defence system. Microbiology 155, 733–740.

Nakayama, K., Sei, N., Oshima, Y., Tashiro, K., Shimasaki, Y., Honjo, T., 2008. Alteration of gene expression profiles in the brain of Japanese medaka (*Oryzias latipes*) exposed to KC-400 or PCB126. Mar. Pollut. Bull. 57, 460–466. https://doi.org/10.1016/j.marpolbul.2008.02.020

Nassef, M., Kato-Unoki, Y., Furuta, T., Nakayama, K., Satone, H., Shimasaki, Y., Honjo, T., Oshima, Y., 2011a. Molecular cloning, sequencing, and gene expression analysis of tributyltin-binding protein type 1 in Japanese medaka fish, *Oryzias latipes*. Zoolog. Sci. 28, 281–285. https://doi.org/10.2108/zsj.28.281

Nassef, M., Tawaratsumita, T., Oba, Y., Satone, H., Nakayama, K., Shimasaki, Y., Honjo, T., Oshima, Y., 2011b. Induction of tributyltin-binding protein type 2 in Japanese flounder, *Paralichthys olivaceus*, by exposure to tributyltin-d27. Mar. Pollut. Bull. 62, 412–414. https://doi.org/10.1016/j.marpolbul.2010.12.005

Nogueira, P., Lourenço, J., Rodriguez, E., Pacheco, M., Santos, C., Rotchell, J.M., Mendo, S., 2009. Transcript profiling and DNA damage in the European eel (*Anguilla anguilla* L.) exposed to 7,12-dimethylbenz[a]anthracene. Aquat. Toxicol. 94, 123–130. https://doi.org/10.1016/j.aquatox.2009.06.007

Oba, Y., Shimasaki, Y., Oshima, Y., Satone, H., Kitano, T., Nakao, M., Kawabata, S.-I., Honjo, T., 2007. Purification and characterization of tributyltin-binding protein type 2 from plasma of Japanese flounder, *Paralichthys olivaceus*. J. Biochem. 142, 229–238. https://doi.org/10.1093/jb/mvm119

Oba, Y., Yamauchi, A., Hashiguchi, Y., Satone, H., Miki, S., Nassef, M., Shimasaki, Y., Kitano, T., Nakao, M., Kawabata, S., Honjo, T., Oshima, Y., 2011. Purification and characterization of tributyltin-binding protein of tiger puffer, *Takifugu rubripes*. Comp. Biochem. Physiol. Part C Toxicol. Pharmacol. 153, 17–23. https://doi.org/10.1016/j.cbpc.2010.07.009

Robinson, N., Baranski, M., Mahapatra, K.D., Saha, J.N., Das, S., Mishra, J., Das, P., Kent, M., Arnyasi, M., Sahoo, P.K., 2014. A linkage map of transcribed single nucleotide polymorphisms in rohu (*Labeo rohita*) and QTL associated with resistance to Aeromonas hydrophila. BMC Genomics 15, 541. https://doi.org/10.1186/1471-2164-15-541

Satone, H., Lee, J.M., Oba, Y., Kusakabe, T., Akahoshi, E., Miki, S., Suzuki, N., Sasayama, Y., Nassef, M., Shimasaki, Y., Kawabata, S., Honjo, T., Oshima, Y., 2011. Tributyltin-binding protein type 1, a lipocalin, prevents inhibition of osteoblastic activity by tributyltin in fish scales. Aquat. Toxicol. 103, 79–84. https://doi.org/10.1016/j.aquatox.2011.02.009

Satone, H., Nonaka, S., Lee, J.M., Shimasaki, Y., Kusakabe, T., Kawabata, S., Oshima, Y., 2017. Tetrodotoxin-and tributyltin-binding abilities of recombinant pufferfish saxitoxin and tetrodotoxin binding proteins of *Takifugu rubripes*. Toxicon 125, 50–52. https://doi.org/10.1016/j.toxicon.2016.11.245

Satone, H., Oshima, Y., Shimasaki, Y., Tawaratsumida, T., Oba, Y., Takahashi, E., Kitano, T., Kawabata, S., Kakuta, Y., Honjo, T., 2008. Tributyltin-binding protein type 1 has a distinctive lipocalin-like structure and is involved in the excretion of tributyltin in Japanese flounder, *Paralichthys olivaceus*. Aquat. Toxicol. 90, 292–299. https://doi.org/10.1016/j.aquatox.2008.08.019

Shimasaki, Y., Oshima, Y., Yokota, Y., Kitano, T., Nakao, M., Kawabata, S., Imada, N., Honjo, T., 2002. Purification and identification of a tributyltin-binding protein from serum of Japanese flounder, *Paralichthys olivaceus*. Environ. Toxicol. Chem. 21, 1229–1235. https://doi.org/10.1002/etc.5620210616

Tian, J., Hu, J., Chen, M., Yin, H., Miao, P., Bai, P., Yin, J., 2017. The use of *mrp1*-deficient (*Danio rerio*) zebrafish embryos to investigate the role of Mrp1 in the toxicity of cadmium chloride and benzo[a]pyrene. Aquat. Toxicol. 186, 123–133. https://doi.org/10.1016/j.aquatox.2017.03.004

Wagner, G.P., Kin, K., Lynch, V.J., 2012. Measurement of mRNA abundance using RNA-seq data: RPKM measure is inconsistent among samples. Theory Biosci. 131, 281–285. https://doi.org/10.1007/s12064-012-0162-3

Wiedenheft, B., Sternberg, S.H., Doudna, J.A., 2012. RNA-guided genetic silencing systems in bacteria and archaea. Nature 482, 331–338. https://doi.org/10.1038/nature10886

Yotsu-Yamashita, M., Nagaoka, Y., Muramoto, K., Cho, Y., Konoki, K., 2018. Pufferfish Saxitoxin and Tetrodotoxin Binding Protein (PSTBP) Analogues in the Blood Plasma of the Pufferfish *Arothron nigropunctatus, A. hispidus, A. manilensis*, and *Chelonodon patoca*. Mar. Drugs 16, 224. https://doi.org/10.3390/md16070224

